# Predicting gene and protein expression levels from DNA and protein sequences with Perceiver

**DOI:** 10.1101/2022.09.21.508821

**Authors:** Matteo Stefanini, Marta Lovino, Rita Cucchiara, Elisa Ficarra

## Abstract

**Background and Objective:** The functions of an organism and its biological processes result from the expression of genes and proteins. Therefore quantifying and predicting mRNA and protein levels is a crucial aspect of scientific research. Concerning the prediction of mRNA levels, the available approaches use the sequence straddling the Transcription Start Site (TSS) as input to neural networks. The State-of-the-art models (e.g., Xpresso and Basenjii) predict mRNA levels exploiting Convolutional (CNN) or Long Short Term Memory (LSTM) Networks. However, CNN prediction depends on convolutional kernel size, and LSTM suffers from capturing long-range dependencies in the sequence. Concerning the prediction of protein levels, as far as we know, there is no model for predicting protein levels by exploiting the gene or protein sequences.

**Methods:** Here, we exploit a new model type (called Perceiver) for mRNA and protein level prediction, exploiting a Transformer-based architecture with an attention module to attend to long-range interactions in the sequences. In addition, the Perceiver model overcomes the quadratic complexity of the standard Transformer architectures. This work’s contributions are 1. DNAPerceiver model to predict mRNA levels from the sequence straddling the TSS; 2. ProteinPerceiver model to predict protein levels from the protein sequence; 3. Protein&DNAPerceiver model to predict protein levels from TSS-straddling and protein sequences.

**Results:** The models are evaluated on cell lines, mice, glioblastoma, and lung cancer tissues. The results show the effectiveness of the Perceiver-type models in predicting mRNA and protein levels.

**Conclusions:** This paper presents a Perceiver architecture for mRNA and protein level prediction. In the future, inserting regulatory and epigenetic information into the model could improve mRNA and protein level predictions. The source code is freely available at https://github.com/MatteoStefanini/DNAPerceiver

**Graphical Abstract:** 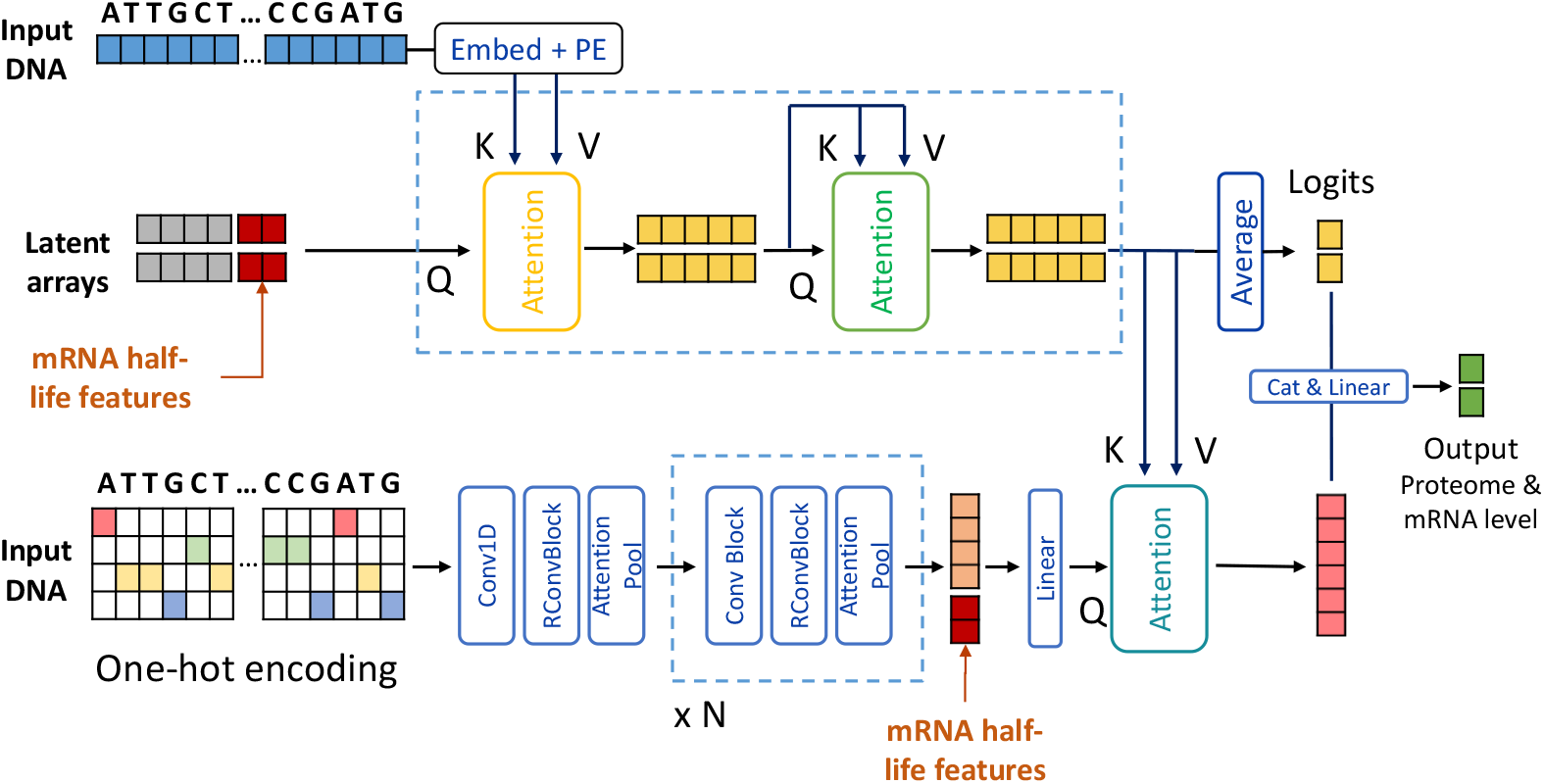

**Highlights:**

- Predicting mRNA and protein levels from DNA and protein sequences is crucial in clinical applications.
- A transformer-based architecture with asymmetric attention (Perceiver) is exploited for mRNA and protein level prediction.
- The Perceiver architecture attends to longer range interactions compared to Transformer, CNN, and LSTM.
- The proposed model achieves state-of-the-art performance for mRNA level prediction.
- To the best of our knowledge, the protein level prediction task is addressed.
- The proposed model is tested on glioblastoma and lung cancer tissues.

## 1. Introduction

Most of the biological processes that regulate the functions of an organism are due to the activity of proteins [1, 2, 3]. In recent decades, the incredible development of sequencing techniques and proteomics quantifications have enabled a systematic analysis of the activity level of thousands of genes and proteins [4, 5]. In addition, it is known that many regulatory and epigenomic processes regulate the expression of mRNAs and proteins [6, 7, 8], and the sequence straddling the transcription start site (TSS) has long been investigated to predict the mRNA levels in various tissues. However, the protein level prediction from sequences has yet to be addressed to the best of our knowledge.

In recent years, deep learning techniques spread in health applications[9, 10, 11, 12] [13, 14] and previous works focused on mRNA level prediction from TSS-straddling sequences [15, 16, 17]. In particular, Convolutional Neural Networks have been adopted to deal with the sequential nature of the DNA[15, 16, 17, 18, 19].

Specifically, Basenjii[15] applies convolutional layers followed by dilated convolutions to share information across large distances in the gene sequences. Dilated convolutions have a wider filter created by inserting spaces in the filter elements. Those gaps exponentially increase the receptive field width, thus taking into account longer dependencies in the sequences.

Similarly, Expecto[16] applies convolutional layers to extract features from the sequences using a predefined window’s size. Each window yields a set of features stacked together in a high-dimensionality feature vector. Spatial transformations are then applied to reduce feature vector dimensionality to output mRNA levels.

On the same line, Xpresso[17] introduced a deep convolutional model composed of two sequential convolutional and max-pooling layers followed by two fully connected layers, demonstrating that a localized region around the transcription start site captures the most relevant information for mRNA level prediction.

Although convolutions represent an effective way to deal with gene sequences, they have some significant limitations that hinder their representational power. Above all, the locality nature of convolutions limits the information propagation in the network among distal elements, requiring many successive layers to expand the receptive field and thus not allowing to capture of long-range relationships and dependencies in sequence elements[20].

In 2017 the attention mechanism revolutionized sequence processing, achieving outstanding performance in capturing long-range dependencies due to each token’s global interaction in the input sequence (so-called self-attention), extracting global information directly from the first layer [20]. However, the self-attention operator has a quadratic complexity *O*(*n*^2^), making the prediction unfeasible for long sequences.

The Enformer model[18] firstly applies self-attention to genomic data, capturing wide-ranging relationships and improving mRNA level prediction. However, to keep the computation feasible, the model is composed of a first convolutional step that extracts local features that are then applied to self-attention layers to capture long-range interactions.

Our method, instead, is based on the Perceiver architecture[21], which allows for asymmetric attention between inputs and learnable query vectors, therefore expanding its capabilities to attend longer sequences directly on the raw data without an initial convolutional step. The advantage of the Perceiver architecture is not limited to the computational aspects. The regulatory parts of a gene (e.g., enhancer and silencer) can be at a considerable distance from the gene region on which they act. Unlike CNN and LSTM, these long-range interactions are modeled in the Perceiver architecture, allowing a better mRNA level prediction.

In this work, we present three models, all based on the Perceiver architecture: DNAPerceiver, ProteinPerceiver, and DNA&ProteinPerceiver. DNAPerceiver predicts the mRNA and protein levels from the TSS-straddling sequence, and its performances are directly compared with competitor models on various datasets. ProteinPerceiver and DNA&ProteinPerceiver instead predict the protein levels from the protein sequence (ProteinPerceiver) and the combination of TSS-straddling and protein sequences (DNA&ProteinPerceiver), respectively. The latter two models were evaluated under different experimental conditions. However, due to the task’s novelty, it is impossible to report comparisons with models in the literature. Below, the Materials and Method section contains the technical details of the models developed and the data used. Subsequently, the Results and Discussion sections report the results obtained. Finally, the Conclusion section summarizes the main contributions of this work.

## 2. Materials and Methods

In order to predict mRNA and protein levels, human protein-coding genes were selected, and their TSS-straddling and protein sequences were obtained (see details in the Dataset section). Then, three models based on a Perceiver architecture were implemented: DNAPerceiver, ProteinPerceiver, and DNA&ProteinPerceiver. DNAPerceiver predicts mRNA levels from the TSS-straddling gene sequence. ProteinPerceiver and DNA&ProteinPerceiver instead predict protein levels from the protein sequence and the combination of the TSS-straddling and protein sequences, respectively. Although each model differs in the prediction task, the general structure is very similar. First, each model receives a sequence representing the TSS-straddling or the protein sequence as input. Then, the Perceiver model encodes and processes the sequence, and a discrete number is outputted for each sequence. This number represents the samples’ average amount of mRNA or protein levels. The greater the number, the greater the amount of molecule (mRNA or protein) circulating. Therefore the main difference between the three Perceiver architectures consists of the input data: TSS-straddling sequences for DNAperceiver; protein sequences for ProteinPerceiver, and TSS-straddling and protein sequences for DNA&ProteinPerceiver architecture.

### 2.1. Datasets

We evaluate our models using different settings, depending on the desired predicted output and the input sequences. There are two input types: inputDNA and inputPROT.

InputDNA consists of the sequence of human protein-coding genes straddling the transcription start site (TSS). The sequence upstream of the TSS contains the gene’s promoter, while the sequence downstream of the TSS contains the exons and introns of the gene. InputDNA sequences are taken from the Xpresso publication [17] due to its particular data curation. Indeed, in this dataset, the TSS positions were accurately revised by Xpresso’s authors exploiting Cap Analysis Gene Expression (CAGE) experiments, a method to measure the actual TSS location. Specifically, it comprises 18377 genes split into 16377 genes for training, 1000 for validation, and 1000 for the test. In addition, the maximum length of the TSS-straddling sequence of a gene is set to 20000 base pairs. Xpresso input also comes with half-life features, which contain general information about the gene (e.g., gene length, number of introns). Therefore, whenever we use InputDNA sequences, we also include half-life features as additional input to our models at different network points, as explained in the architecture section.

InputPROT, on the other hand, consists of protein sequences. Therefore, the promoter region and all non-coding parts of a gene are not included in the inputPROT sequence. All protein sequences were obtained from Uniprot database [22], processed with Biopython library [23], and intersected with Xpresso’s list of protein-coding genes.

As for the labels, we used four conditions for predicting mRNA levels (labelExprMouse, labelExprHuman, labelGeneGlio, labelGeneLung) and two conditions for predicting protein levels (labelProtGlio and labelProtLung). LabelExprMouse and labelExprHuman come from the Xpresso publication, containing the mean mRNA levels of mouse and human samples, respectively. These labels were obtained in the biologically controlled context of cell lines, and therefore the prediction task is limited. To evaluate the predictive capabilities of the models on high throughput multi-omics human data from clinical studies, we selected mRNA and protein levels on patients with glioblastoma [24] and lung cancer [25]. LabelGeneGlio and labelGeneLung contain the labels of the mediated mRNA values for glioblastoma and lung cancer, respectively. The same procedure has been applied to obtain the mediated protein levels for the same patients, named LabelProtGlio and labelProtLung, respectively [24, 25].

Given the scarcity of data, except for Xpresso comparisons, we adopt the K-Fold validation setting and average the results across the folds. We set the number of folds K to 10.

### 2.2. Metric

To measure the effectiveness of our methods, we compute the variance explained *r*^2^, also known as the coefficient of determination: 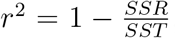, where SSR stands for Sum Squared Regression (the sum of the residuals -actual values minus predicted value-squared) and SST for Total Sum of Squares (the sum of the distance the data is away from the mean all squared). This coefficient is the most widely adopted metric for mRNA level prediction, ranging from 0 to 1. When it is 0, the model makes a prediction no better than random, while when it is 1 the model perfectly predicts the actual labels.

### 2.3. DNAPerceiver architecture

As stated above, various models in the literature focused on predicting mRNA levels from the TSS-straddling sequence. This work aims to reveal if mRNA levels can be explained by the TSS-straddling sequence alone. All predictive models do not use the whole gene sequence as input but only the portion straddling TSS, which involves numerous regulatory and transcriptional processes. In particular, the region preceding the TSS contains the promoter, a region targeted explicitly by transcription factors, elements responsible for the final quantity of mRNA produced. The data used in this model are inputDNA as input and labelExprMouse, labelExprHuman, labelGeneGlio, and labelGeneLung as output.

Figure 1 shows the architecture of the DNAPerceiver. The model is composed of two distinct flows: one with asymmetric attention as in the original Perceiver model [21], and another with a convolutional step inspired by the Enformer model [18]. The asymmetric attention reduces the complexity of the attention from *O*(*n*^2^) to *O*(*n × m*) where *n* is the length of the input sequence, and *m* is a hyperparameter defying the latent space dimensionality. The model can attend to long sequences and condense their semantic information within a tight latent space. The convolutional step extracts another representation of the same DNA sequence and is then used to query the latent space in the final decoding stage.

**Figure 1:**
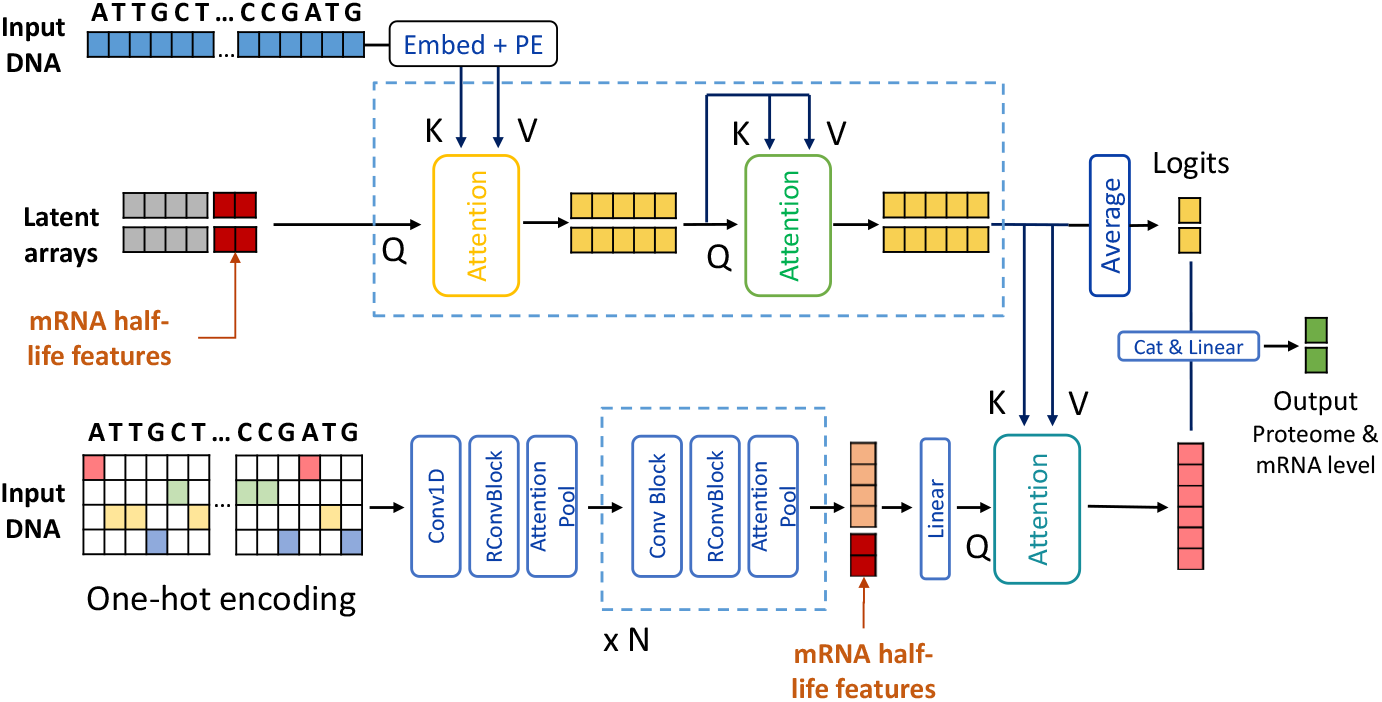
DNAPerceiver architecture. It is based on the Perceiver IO model [26]. The upper flow represents the asymmetric attention that distills the sequence in a smaller latent space, where learnable arrays attend to all the input sequences and refine their representations with self-attention and feed-forward networks. The lower flow depicts the decoding stage of the Perceiver IO, where instead of using learnable vectors like in the original model, we use, as the final query, the same sequence processed by a convolutional pipeline inspired by the Enformer model [18]. In this figure, Q,K,V stands for Query, Keys and Values as in typical Transformer architecture, PE is the Positional Encoding, Conv1D is a 1-dimensional Convolution and RConvBlock is a 1D Convolution with a residual connection. The first Convolutional layer is applied to the one-hot encoded version of the sequence, as all previous model of literature, while the upper part of the model embeds the one-hot vectors into learnable embedding vectors through linear projections, as typical Transformer architecture requires.

Therefore, while our model still leverages a convolutional step, it takes more advantage of the recent advancements of attentive architectures, *i.e*. the Perceiver, that have originated from the original Transformer model. Transformers are a class of deep learning models, first introduced by Vaswani *et al*. [20], that attained substantial breakthroughs in natural language processing and computer vision. Specifically, they consist of attention blocks that aggregate information from the entire input sequence by computing a weighted sum across the representations of all other tokens for each sequence token. Since each token directly attends to all other positions in the sequence, they allow for a much better information flow between distal elements, in contrast with convolutional layers, which may require many successive layers to increase the receptive field [20].

These methods were recently applied to model mRNA sequences. However, given the quadratic complexity of attention *O*(*n*^2^), the length of the input can explode quadratically, rendering it infeasible to encode sequences of more than a few thousand letters. For this reason, our method is based on the Perceiver [21], a model that builds upon Transformers but scales to hundreds of thousands of inputs, as it leverages an asymmetric attention mechanism to distill inputs into a tight latent bottleneck iteratively. Then, the latent arrays go through self-attention blocks to refine their representation and potentially other asymmetric attention layers before getting averaged to obtain the logits for the task at hand.

Specifically, we use the Perceiver IO [26], which improved the decoder capabilities of the model by adding a final decoding stage. This stage acts as a query on the latent arrays, allowing the model to produce outputs of arbitrary size and semantics, and deal with diverse domains without sacrificing the benefits of deep, domain-agnostic processing.

In our implementation, however, we introduce substantial modifications concerning the Perceiver IO architecture. Firstly, instead of learning a different set of output arrays for the decoding stage, we use the same InputDNA sequence after being processed by a Convolutional step. This step consists of multiple Conv layers, Residual connections, and Attention Pooling layers inspired by the Enformer model [18]. Secondly, another difference is that our model in the decoding stage also considers the processed latent arrays by applying a final head that computes their average and uses them as final logits. The processing is similar to that of the original Perceiver model. However, in our case, it is fused with the final decoding mechanism proposed by the Perceiver IO.

Hence, in our architecture, the TSS-straddling or the protein sequence given in input is processed twofold: as learnable vectors for the perceiver flow, where the asymmetric attention is applied with the latent arrays, and as one-hot encoding vectors fed to the convolutional step. After embedding the input letters, we also add a learnable Positional Encoding, initialized with a sinusoidal function as in the original Transformer model to deal with positions in the asymmetric attention.

Latent arrays are initialized with random numbers from a normal distribution with mean 0 and variance 1, while the inputDNA is represented with one-hot encoding vectors, applied to the Convolutional step, and linearly projected into embedding vectors for the attention path. Moreover, mRNA half-life features are injected in both flows: they are appended to the latent arrays and the convolutional step in the final feature representations.

In the DNAPerceiver configuration, we predict the mRNA level for both human cell-line and mouse data, and we evaluate our model on the Xpresso dataset, comparing it with other similar methods and Xpresso itself. Further, we applied the DNAPerceiver model to predict mRNA and protein levels in human high throughput sequencing data. As discussed in the Dataset section, we take the protein labels from two real-human datasets, ending with 10529 pairs of labels for Lung cancer (labelProtLung) and 10280 pairs of labels for glioblastoma (labelProtGlio). In this configuration, we output two predictions for labelGene and labelProt, mRNA and protein levels, respectively, for both datasets.

To represent the A, T, C, and G letters, we use one-hot vectors, and for the perceiver flow, we linearly project them to learnable vectors of dimensionality 32. In addition, we add the letter P as padding. To represent letter positions, we employ learnable positional encodings initialized in a standard sinusoidal fashion[20]. We use 128 latent learnable arrays with a dimensionality of 128 each, constituting the dimensionality of the following self-attention layers. The number of heads in asynchronous attention is set to 1, while self-attention is set to 8. The attention over the input is computed by considering only valid letters and masking the rest. Feed-forward layers have a dimensionality of 256 and GELU nonlinearity. The depth of the Perceiver, the number of layers of asynchronous attention followed by self-attention, is set to 1. For the convolutional query flow, we adopt a similar strategy as Enformer[18] using the first layer of Conv1D with a kernel size of 15, channel dimensionality of 64 and Attention Pooling with a pooling size of 12. Subsequent convolutional layers, forming the Conv tower, have a kernel size of 5 and attention pooling size of 6. Each Conv layer applies GELU nonlinearity and is followed by a residual connection. The length of the InputDNA sequence is set to 10500, taking the majority part from the promoter side and less from the actual gene, specifically considering 7000 base pairs before the TSS and 3500 after the TSS. We apply dropout throughout the model before each linear projection and attention layer, with a keep probability of 0.8. We train our model using ADAM optimizer[27], a batch size of 128, and we follow the learning rate scheduling strategy of [20] with a warmup equal to 8000 iterations. We apply a weight decay of 0.2 and an early stopping strategy to avoid overfitting. We found it helpful to use the Tanh activation for our final predicted scores only in this configuration and when applied to Xpresso mRNA levels. In the end, we weighted the loss contribution using a weight of 10 for the mRNA.

### 2.4. ProteinPerceiver architecture

Although the mRNA level prediction task is debated within the scientific community, to the best of our knowledge, there are no publicly available models for protein level prediction using protein sequences. In the last decade, the quantification of mRNA levels has been available in large quantities. Instead, extracting and quantifying proteins is more recent and less mature than mRNA extraction and quantification techniques. Protein quantification is what scientists are most interested in biologically. However, these techniques are currently more expensive, limiting data availability. Moreover, the mRNA level quantification can evaluate more than 20000 protein-coding genes versus approximately 2 to 8 thousand proteins for protein quantification. The ProteinPerceiver model aims to measure how much protein levels depend on the protein sequence. Unfortunately, given the experiments’ novelty, no comparison model in the literature is available. The input consists of inputProtein and labelProtGlio and labelProtLung as output.

We match protein sequences and proteomics labels available, assembling a total of 10430 protein sequences for Glioblastoma and 10699 for Lung Carcinoma with corresponding proteomic labels.

Differently from the DNAPerceiver setting, here we only adjust the dropout keep probability to 0.7 and the attention pooling size in the convolutional query to 10 in the first layer and 5 in the following ones. Moreover, we set the maximum length of the protein sequence to 6000 and the final weight of the MSE loss to 100 for Lung data and 3000 for Glioblastoma data. We optimize the model using Lamb[28], a learning rate of 0.0005, and a Cosine Annealing schedule strategy with 8000 steps of warmup.

### 2.5. Protein&DNAPerceiver architecture

The ProteinPerceiver model receives the protein sequence as input to predict protein levels. However, the protein level is determined by the protein sequence and by regulatory, transcriptional, and epigenetic factors. Although considering all regulatory processes is not straightforward, in this paper, we have evaluated the combined effect of the protein and the TSS-straddling sequence to predict the protein levels. The model simultaneously uses inputDNA and inputProtein and outputs labelProtGlio and labelProtLung.

TSS-straddling and protein sequences are matched together when both are available from the Xpresso dataset[17] and the protein sequence dataset, ending up with a total of 9815 triplets gene-protein-labels for Lung Carcinoma and 9534 triplets for Glioblastoma.

Explicitly, our model deals with two different input sequences, one for the Perceiver flow and one for the Convolutional query. In addition, we investigated the use of the input in an alternate manner: when the TSS-straddling sequence is in input to the Perceiver, we use the protein sequence as a query, and vice versa, with protein as the perceiver input, we use the TSS-straddling for the query computation. The Results section shows that the best version differs depending on the data and the prediction. The maximum length of the protein sequence is set to 6000, while the DNA sequence length is set to 8000. If not specified, we kept the same hyperparameters of the DNAPerceiver configuration.

A summary of the architecture names, prediction tasks, input, and outputs is reported in Table 1.

**Table 1:** Summary of the different configurations of our model depending on the prediction task and the input-output setting.

| Model | Tasks | Input | Output |
| --- | --- | --- | --- |
| DNAPerceiver | mRNA levels | InputDNA | labelGene |
| DNAPerceiver | mRNA&protein levels | InputDNA | labelGene&labelProt |
| ProteinPerceiver | protein levels | InputProt | labelProt |
| DNA&ProteinPerceiver | protein levels | InputDNA&InputProt | labelProt |

## 3. Results

This section discusses the results obtained and the comparison with the state-of-the-art approaches.

### 3.1 Results on mRNA level prediction using Xpresso’s labels

In this setting, DNAPerceiver was trained on the Xpresso sequences and their labels, aiming to predict the mRNA level, both from mouse and human organisms (labelExprMouse and labelExprHuman). We follow the split of the original dataset[17], thus obtaining 16377 genes for training and 1000 genes for both validation and test set. As shown in Table 2, DNAPerceiver performs better than the Xpresso method in terms of *r*^2^ in human and mouse data. In human cell-line data, it reaches an *r*^2^ of 0.62, which, compared to the 0.59 of the Xpresso model, gains 0.03 points of *r*^2^.

**Table 2:**
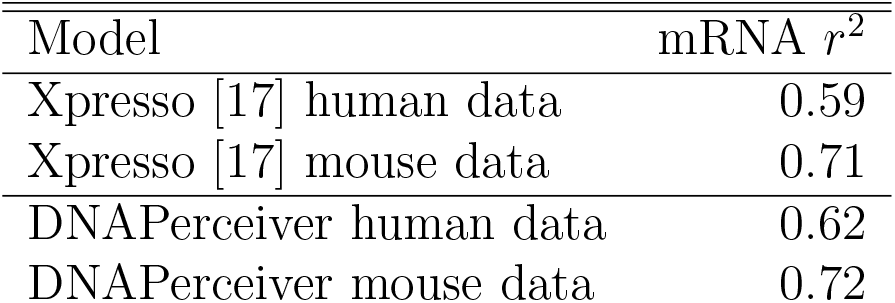
Results on the test set of Xpresso dataset in predicting mRNA levels of cell-line data. The input is the InputDNA sequence, and the output is the mRNA level, expressed with the coefficient of determination *r*^2^.

The basenji method has a similar mRNA level prediction task to the one presented in this work. However, a direct comparison cannot be made as Basenji uses Cap Analysis Gene Expression (CAGE) input data which are not available for our dataset (Xpresso’s dataset released the sequences but not the CAGE information). However, under his experimental conditions, Basenji reaches a Pearson correlation coefficient ranging from 0.138 to 0.777, depending on the genes considered. These values would translate into a coefficient of determination *r*^2^ between 0.019 and 0.604. In this context, the DNAPerceiver model gets consistent results.

### 3.2. Results on mRNA and protein levels

In this configuration, DNAPerceiver is trained on inputDNA (Xpresso sequences) and predicts the labels from the Lung and Glioblastoma datasets for mRNA and protein levels (labelGeneLung, labelGeneGlio, labelProtLung, and labelProtGlio). Table3 reports the results. High-throughput sequencing data from human tissues is much more complex than data obtained from cell lines. Indeed, the cell lines are systematically obtained in the laboratory to have a controlled context and genetic variability as small as possible between the cells. By contrast, the sequencing data from tissues (tumor tissues, too) has a high genetic variability as a multiplicity of regulatory factors between cells and tissues are present. Given the noisy nature of high throughput sequencing data, its mRNA level prediction is not comparable to that of a cell line culture, but it reaches 0.181 of *r*^2^. Furthermore, our focus is to predict the protein level using only the InputDNA sequence. As a result, our model can predict the protein levels achieving 0.161 of *r*^2^, demonstrating its capability to perceive the direct connection between the InputDNA sequence and its corresponding protein level.

**Table 3:** Results on Lung and Glioblastoma data in predicting mRNA and protein level. The input is inputDNA, taken from Xpresso[17] publication, while predicted labels for mRNA and protein levels are labelGeneLung, labelGeneGlio, labelProtLung, and label-ProtGlio. Results are the average of the k-fold validation method with k equal to 10.

| Model | mRNA $r^2$ | proteomics $r^2$ |
| --- | --- | --- |
| DNAPerceiver Lung | 0.181 | 0.161 |
| DNAPerceiver Glioblastoma | 0.150 | 0.026 |

### 3.3. Results on protein level using protein sequence as input

Table4 reports the result of our model applied to protein sequence as input. In this configuration of our model, named ProteinPerceiver, the input consists of the protein sequences, as explained in the Datasets subsection in the Material and methods Section. The aim is to predict the protein level given the protein sequence, a peculiarly complex task, as discussed later in the next section. The obtained outcome varies depending on the data: for Lung data, we found that predicting protein levels from the protein sequence is more complex, achieving a *r*^2^ of 0.085, comparing the 0.161 obtained from the InputDNA. Nonetheless, for Glioblastoma data, our ProteinPerceiver can score a *r*^2^ of 0.028 for protein levels, which is slightly better compared to 0.026 obtained by the DNAPerceiver.

**Table 4:** Results in predicting protein levels from the protein sequence. The input is InputPROT, while predicted labels for protein levels are labelProtLung and labelProtGlio. Results are the average of the k-fold validation method with k equal to 10.

| Model | proteomics $r^2$ |
| --- | --- |
| ProteinPerceiver Lung | 0.085 |
| ProteinPerceiver Glioblastoma | 0.028 |

Despite the impact of data quality and prediction task complexity on the results, our model can still capture a part of the relationship between the protein sequence and its corresponding protein level.

### 3.4. Results on protein levels using TSS-straddling and protein sequences as input

We wanted to investigate further the model’s capabilities with a peculiar configuration, in which we give as input both the TSS-straddling (InputDNA) and the protein sequence. Our model’s design facilitates this accomplishment, already treating the input in a double structure: one for the perceiver flow and one for the convolutional query. Therefore, the protein sequence was input to the perceiver and the InputDNA to the convolutional query and vice-versa. We report the results in Table5. In this configuration, performances also depend on the specific data: for Lung data, surprisingly, the use of both inputs does not improve the total performances of the model, reaching 0.141 of *r*^2^ compared to the 0.161 obtained using only InputDNA sequence. On the contrary, using both inputs slightly improves the results on Glioblastoma data, achieving 0.031 of *r*^2^.

**Table 5:** Results in predicting protein levels from both the DNA sequence and the protein sequence used as inputs. The input is InputPROT and inputDNA, and the predicted labels for protein expression are labelProtLung and labelProtGlio. Results are the average of the k-fold validation method with k equal to 10.

| Model | proteomics $r^2$ |
| --- | --- |
| Protein&DNA Perceiver Lung | 0.141 |
| Protein&DNA Perceiver Glioblastoma | 0.031 |

## 4. Discussion

In regards to predicting mRNA levels from the sequence straddling the TSS (thus including part of the promoter and part of the gene), DNAPerceiver shows results superior to Xpresso in the case of the human cell lines and murine samples. Unlike the Xpresso model, the DNAPerceiver model exploits the self-attention mechanism to predict the mRNA levels. Having the same input sequence size and output levels as Xpresso, the DNAPerceiver model achieves superior results since long-range interactions between the most distant regions of the promoter and the gene sequence are fully exploited in the model and not limited by the size of the convolutional kernel. Moreover, as can be expected, the prediction of mRNA levels in cell lines achieves better results than mRNA level prediction in tumor samples. This aspect could be explained by the different boundary conditions of the two situations. In the first case, the mRNA expression is controlled to ensure the reproducibility and stability of the cell lines. In the second case, the intrinsic samples’ variability cannot be limited and pathological conditions profoundly alter the biological context. Besides the improvement in mRNA level prediction, the main novelty of this work is the prediction of protein levels from the TSS-straddling and protein sequences. This aspect is doubly challenging: 1. Protein extraction and quantification techniques have emerged recently, so data availability still needs to be improved compared to mRNA datasets; 2. The protein sequence has target regions for post-translation regulators; however, the promoter region is not used as input in the ProteinPerceiver model. It is noted that the prediction of protein levels is considerably lower than mRNA ones, whether the prediction exploits the TSS-straddling or the protein sequence. The complexity of the problem can explain this phenomenon. The protein level is influenced by notable post-transcriptional and post-translational regulatory phenomena (e.g., ubiquitination), which are not fed to the models. Moreover, the TSS-straddling sequences (composed of the promoter and a part of the gene) have a greater predictive power of the protein level than the protein sequences. This behavior could depend on the presence of the promoter. Indeed, the promoter is the region that favors the expression regulation (both of genes and, therefore, of proteins), and it is responsible for interacting with transcription factors. When the model is trained simultaneously with the TSS-straddling and the protein sequence, the predictive power of protein level increases; however, it remains lower than the prediction of protein levels using only the TSS-straddling sequence. In this sense, the TSS-straddling sequence seems more informative than the protein one.

## 5. Conclusions

Various papers have addressed mRNA level prediction in the literature, which mainly includes convolutional or long short-term memory networks. This work presents three Perceiver-type architectures for predicting mRNA levels on cell lines and high-throughput human samples. Furthermore, a novel task is introduced, presenting the prediction of protein levels from the TSS-straddling and protein sequences. The results show the advantages of the Perceiver architecture in predicting mRNA levels compared to competitors. On the other hand, protein level prediction benefits more from the TSS-straddling sequence than the protein one. This aspect could be explained by the presence of the promoter region in the TSS-straddling sequence.

The Perceiver architecture benefits are not limited to the computational aspects. Since the regulatory parts of a gene (e.g., enhancer and silencer) can be at a considerable distance from the TSS, these regions can be directly attended by Perceiver models. Furthermore, unlike CNN and LSTM, long-range interactions can be exploited in the Perceiver architecture, allowing a better prediction.

Although various experimental conditions have been considered, other biological post-transcriptional and post-translation regulations can be included in the models to improve prediction.

## References

[1] F. Crick, L. Barnett, S. Brenner, R. J. Watts-Tobin, et al., General nature of the genetic code for proteins, Nature (1961).

[2] F. Crick, Central dogma of molecular biology, Nature 227 (5258) (1970) 561–563.

[3] A. Wada, H. Nakamura, Nature of the charge distribution in proteins, Nature 293 (5835) (1981) 757–758.

[4] F. Zhang, W. Ge, G. Ruan, X. Cai, T. Guo, Data-independent acquisition mass spectrometry-based proteomics and software tools: a glimpse in 2020, Proteomics 20 (17-18) (2020) 1900276.

[5] E. Phizicky, P. I. Bastiaens, H. Zhu, M. Snyder, S. Fields, Protein analysis on a proteomic scale, Nature 422 (6928) (2003) 208–215.

[6] E. Jablonka, M. J. Lamb, The changing concept of epigenetics, Annals of the New York Academy of Sciences 981 (1) (2002) 82–96.

[7] A. Bird, Perceptions of epigenetics, Nature 447 (7143) (2007) 396.

[8] M. Esteller, Epigenetics in cancer, New England Journal of Medicine 358 (11) (2008) 1148–1159.

[9] A. Mascolini, S. Puzzo, G. Incatasciato, F. Ponzio, E. Ficarra, S. Di Cataldo, A novel proof-of-concept framework for the exploitation of convnets on whole slide images, in: Progresses in Artificial Intelligence and Neural Systems, Springer, 2021, pp. 125–136.

[10] S. Allegretti, F. Bolelli, F. Pollastri, S. Longhitano, G. Pellacani, C. Grana, Supporting Skin Lesion Diagnosis with Content-Based Image Retrieval, in: 2020 25th International Conference on Pattern Recognition (ICPR), IEEE, 2021, pp. 8053–8060. doi:10.1109/ICPR48806.2021.9412419.

[11] F. Bolelli, S. Allegretti, C. Grana, One DAG to Rule Them All, IEEE Transactions on Pattern Analysis and Machine Intelligence 44 (7) (2021) 3647–3658. doi:https://doi.org/10.1109/TPAMI.2021.3055337.

[12] M. Cancilla, L. Canalini, F. Bolelli, S. Allegretti, S. Carrión, R. Paredes, J. A. Gómez, S. Leo, M. E. Piras, L. Pireddu, A. Badouh, S. Marco-Sola, L. Alvarez, M. Moreto, C. Grana, The DeepHealth Toolkit: A Unified Framework to Boost Biomedical Applications, in: 2020 25th International Conference on Pattern Recognition (ICPR), IEEE, 2021, pp. 9881–9888.

[13] A. Khan, Z. Rehman, H. F. Hashmi, A. A. Khan, M. Junaid, A. M. Sayaf, S. S. Ali, F. U. Hassan, W. Heng, D.-Q. Wei, An integrated systems biology and network-based approaches to identify novel biomarkers in breast cancer cell lines using gene expression data, Interdisciplinary Sciences: Computational Life Sciences 12 (2) (2020) 155–168.

[14] W. Zhang, X. Xue, C. Xie, Y. Li, J. Liu, H. Chen, G. Li, Cegso: boosting essential proteins prediction by integrating protein complex, gene expression, gene ontology, subcellular localization and orthology information, Interdisciplinary Sciences: Computational Life Sciences 13 (3) (2021) 349–361.

[15] D. R. Kelley, Y. A. Reshef, M. Bileschi, D. Belanger, C. Y. McLean, J. Snoek, Sequential regulatory activity prediction across chromosomes with convolutional neural networks, Genome research 28 (5) (2018) 739–750.

[16] J. Zhou, C. L. Theesfeld, K. Yao, K. M. Chen, A. K. Wong, O. G. Troyanskaya, Deep learning sequence-based ab initio prediction of variant effects on expression and disease risk, Nature genetics 50 (8) (2018) 1171–1179.

[17] V. Agarwal, J. Shendure, Predicting mrna abundance directly from genomic sequence using deep convolutional neural networks, Cell reports 31 (7) (2020) 107663.

[18] Ž. Avsec, V. Agarwal, D. Visentin, J. R. Ledsam, A. Grabska-Barwinska, K. R. Taylor, Y. Assael, J. Jumper, P. Kohli, D. R. Kelley, Effective gene expression prediction from sequence by integrating long-range interactions, Nature methods 18 (10) (2021) 1196–1203.

[19] V. Pipoli, M. Cappelli, A. Palladini, C. Peluso, M. Lovino, E. Ficarra, Predicting gene expression levels from dna sequences and posttranscriptional information with transformers, Computer Methods and Programs in Biomedicine (2022) 107035.

[20] A. Vaswani, N. Shazeer, N. Parmar, J. Uszkoreit, L. Jones, A. N. Gomez, L-. Kaiser, I. Polosukhin, Attention is all you need, Advances in neural information processing systems 30 (2017).

[21] A. Jaegle, F. Gimeno, A. Brock, O. Vinyals, A. Zisserman, J. Carreira, Perceiver: General perception with iterative attention, in: International Conference on Machine Learning, PMLR, 2021, pp. 4651–4664.

[22] U. Consortium, Uniprot: a worldwide hub of protein knowledge, Nucleic acids research 47 (D1) (2019) D506–D515.

[23] P. J. Cock, T. Antao, J. T. Chang, B. A. Chapman, C. J. Cox, A. Dalke Friedberg, T. Hamelryck, F. Kauff, B. Wilczynski, et al., Biopython: freely available python tools for computational molecular biology and bioinformatics, Bioinformatics 25 (11) (2009) 1422–1423.

[24] L.-B. Wang, A. Karpova, M. A. Gritsenko, J. E. Kyle, S. Cao, Y. Li, D. Rykunov, A. Colaprico, J. H. Rothstein, R. Hong, et al., Proteogenomic and metabolomic characterization of human glioblastoma, Cancer Cell 39 (4) (2021) 509–528.

[25] S. Satpathy, K. Krug, P. M. J. Beltran, S. R. Savage, F. Petralia, C. Kumar-Sinha, Y. Dou, B. Reva, M. H. Kane, S. C. Avanessian, et al., A proteogenomic portrait of lung squamous cell carcinoma, Cell 184 (16) (2021) 4348–4371.

[26] A. Jaegle, S. Borgeaud, J.-B. Alayrac, C. Doersch, C. Ionescu, D. Ding, S. Koppula, D. Zoran, A. Brock, E. Shelhamer, et al., Perceiver io: A general architecture for structured inputs & outputs, arXiv preprint 2107.14795 (2021).

[27] J. Zhang, S. P. Karimireddy, A. Veit, S. Kim, S. J. Reddi, S. Kumar, S. Sra, Why adam beats sgd for attention models (2019). 1912.03194.

[28] Y. You, J. Li, S. Reddi, J. Hseu, S. Kumar, S. Bhojanapalli, X. Song, J. Demmel, K. Keutzer, C.-J. Hsieh, Large batch optimization for deep learning: Training bert in 76 minutes, arXiv preprint 1904.00962 (2019).

